# A comprehensive anatomical map of the peripheral octopaminergic/tyraminergic system of *Drosophila melanogaster*

**DOI:** 10.1101/368803

**Authors:** Dennis Pauls, Christine Blechschmidt, Felix Frantzmann, Basil el Jundi, Mareike Selcho

**Author notes:** Correspondence: Mareike Selcho, Neurobiology and Genetics, Theodor-Boveri Institute, Biocenter, University of Würzburg, Am Hubland, 97074 Würzburg, Germany.

## Abstract

The modulation of an animal’s behavior through external sensory stimuli, previous experience and its internal state is crucial to survive in a constantly changing environment. In most insects, octopamine (OA) and its precursor tyramine (TA) modulate a variety of physiological processes and behaviors by shifting the organism from a relaxed or dormant condition to a responsive, excited and alerted state. Even though OA/TA neurons of the central brain are described on single cell level in *Drosophila melanogaster,* the periphery was largely omitted from anatomical studies. Given that OA/TA is involved in behaviors like feeding, flying and locomotion, which highly depend on a variety of peripheral organs, it is necessary to study the peripheral connections of these neurons to get a complete picture of the OA/TA circuitry. We here describe the anatomy of this aminergic system in relation to peripheral tissues of the entire fly. OA/TA neurons arborize onto skeletal muscles all over the body and innervate reproductive organs, the heart, the corpora allata, and sensory neurons in the antenna, wings and halteres underlining their relevance in modulating complex behaviors.

## Introduction

The adrenergic system of mammals influences various aspects of the animal’s life. Its transmitters/hormones, adrenaline and noradrenaline, modulate a variety of physiological processes and behaviors. They are secreted into the bloodstream by the adrenal glands in response to stress. In addition, they are synthesized and released by axonal terminals in the central nervous system (CNS) as well as sympathetic fibers of the autonomic nervous system. Adrenaline and noradrenaline have been described as modulators to shift the organism from a relaxed or dormant state to a responsive, excited and alerted state ^1^. Stressful stimuli induce a metabolic and behavioral adaptation, leading to enhanced energy supply, increased muscle performance, increased sensory perception and a matched behavior. This so-called “fight or flight” response can be seen in vertebrates and invertebrates. In insects, the stress response is mediated - among others - by octopamine (OA) and its precursor tyramine (TA) ^2–4^ TA is synthesized from tyrosine by the action of a tyrosine decarboxylase enzyme (Tdc) and functions as an independent neurotransmitter/-modulator as well as the intermediate step in OA synthesis. For this, TA is catalyzed by the tyramine-β-hydroxylase (TβH).

Similar to the vertebrate adrenergic system, OA and TA act through specific G-protein coupled receptors. Besides structural similarities between OA/TA and adrenaline/noradrenaline and the corresponding receptors, functional similarities are illustrated by the action of these transmitters/hormones in the regulation of physiological processes and behaviors. OA and TA are known to modulate muscle performance, glycogenolysis, fat metabolism, heart rate, and respiration in insects (reviewed by: ^5^).

While the role of TA as an independent signaling molecule was underestimated for a long time, OA has been extensively studied and was shown to have effects on almost every organ, sensory modality and behavior in a great variety of insects. The most intensively studied peripheral organs regarding the modulatory role of OA are muscles ^6–10^ Here, OA is thought to not exclusively modulate muscle performance or motor activity. OA rather modulates muscle action according to metabolic and physiological processes, for example by promoting energy mobilization directly from the fat body, or indirectly by promoting the release of adipokinetic homones (AKH) from neuroendocrine cells in the corpora cardiaca (CC, a homolog of the vertebrate anterior pituitary gland and an analog of mammalian pancreatic alpha cells) ^11,12^. In addition to the impact of OA/TA on muscles, fat body and AKH cells, OA is shown to modulate the heart, trachea and air sacs, gut, hemocytes, salivary glands, Malpighian tubules and ovaries in insects, mainly to induce a general stress or arousal state. However, in total OA seems to modulate a vast number of behaviors, which are not necessarily coupled to stress responses. The OA/TA system is shown to also act on i.a. learning and memory, sleep, feeding, flight, locomotion, and aggression ^8,10,12–35^.

As mentioned above, OA and TA act as neurotransmitters and neuromodulators, allowing them to act in a paracrine, endocrine or autocrine fashion. In the fruit fly *Drosophila,* huge efforts were made to describe OA/TA neurons (OANs/TANs) in the brain and ventral nervous system (VNS) down to the single cell level ^8,16,36–40^. Nevertheless, although our knowledge about physiological processes and behaviors modulated by the OA/TA system in the brain is rich, less is known about how OA and TA reach all its target organs and tissues in the periphery (exceptions: reproductive organs ^36,40–43^ and muscles ^8,44–46^).

Here we use the genetically tractable fruit fly *Drosophila melanogaster* to describe the arborizations of *Tdc2-Gal4*-positive, and therefore OANs and TANs in the periphery, as the *Drosophila* Tdc2 gene is expressed neurally ^40^. We found that OANs/TANs are widespread distributed throughout the fly’s body with innervations in the skeletal muscles, reproductive organs, corpora allata, antenna, legs, wings, halteres and the heart. This diverse innervation pattern reflects the modulatory role of OA/TA in many different behaviors and physiological processes. Our results provide, for the very first time, a complete and comprehensive map of the OA/TA circuitry in the entire insect body. This map allows assumptions about the type of OA/TA signaling (paracrine or endocrine) to a specific organ and, at the same time, it provides a deeper understanding to what extend the OA/TA-dependent activity of peripheral organs is altered, for example by genetically manipulating *Tdc2-Gal4-positive* neurons in the brain and VNS.

## Results

The OANs/TANs of the brain and ventral nervous system (VNS) are described in detail even on single cell level in *Drosophila* ^10,37–39,44^. In contrast little is known about their peripheral arborizations. We used the well-characterized *Tdc2-Gal4* line to deepen our knowledge about the OA/TA system in the entire body of *Drosophila* ^38–41,47^. Nearly all *Tdc2-Gal4*-positive cells in the brain are stained by a Tdc2 antibody ^47^. In the VNS all of the *Tdc2-Gal4*-positive cells were labeled by a TβH antibody and therefore have to be Tdc2-positive ^39^. We here expressed myristoylated GFP, enhanced by GFP antibody staining, to label the membranes of *Tdc2-Gal4-positive* neurons from the soma to its fine endings in the periphery. The peripheral organs, tissues and cells are visualized by fluorescent markers for cell bodies (DAPI binds to DNA), muscles (Phalloidin binds F-actin) and antibodies against the synaptic protein Bruchpilot and the cell adhesion molecule Fasciclin 2 (Fig. 1).

**Fig. 1:**
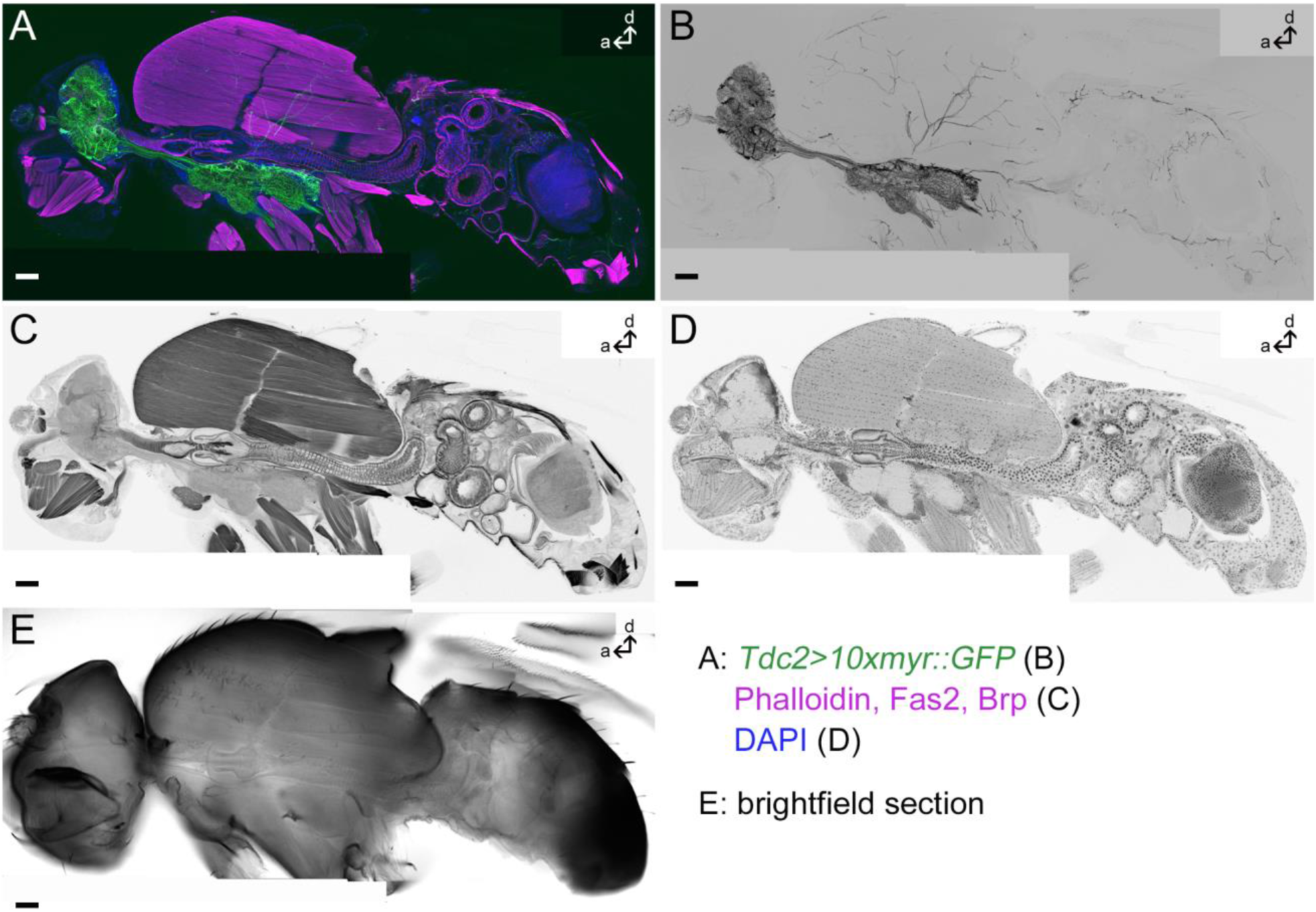
Tdc2-Gal4-positive arborizations in a whole fly section. *Projection of one medial sagittal agarose section of 80μm thickness labeled by anti-GFP to visualize membranes of Tdc2-Gal4-positive neurons (green in A, black in B); Phalloidin, anti-Fasciclin2 (Fas2) and anti-Bruchpilot (Brp) to visualize muscles, cells and synapses, respectively (magenta in A, black in C) and DAPI to mark cell bodies (blue in A, black in D). E: A single optical section showing the bright-field picture. Scale bars = 50 μm*.

### *Tdc2-Gal4-positive* arborizations in the head

*Tdc2-Gal4*-positive neurons (Tdc2N) project through all peripheral nerves of the brain: antennal nerve *(AN),* ocellar nerve (*OCN*), pharyngeal nerve *(PhN)* and accessory PhN *(APhN),* maxillary-labial nerve *(MxLbN),* corpora cardiaca nerve (NCC), and the cervical connective *(CV)* (Fig. 2). Tdc2Ns in the antennal nerve are connected to the antennal lobe and the antennal mechanosensory and motor center (ammc), respectively, and give rise to staining in the pedicle-the Johnston’s organ (JO)-, funiculus and arista of the antenna (Fig. 2A-D). While no cell bodies are visible in the third to fifth segment of the antenna, the JO contains stained cell bodies indicating that the *Tdc2-Gal4* line includes mechanosensory neurons (Fig. 2C).

**Fig. 2:**
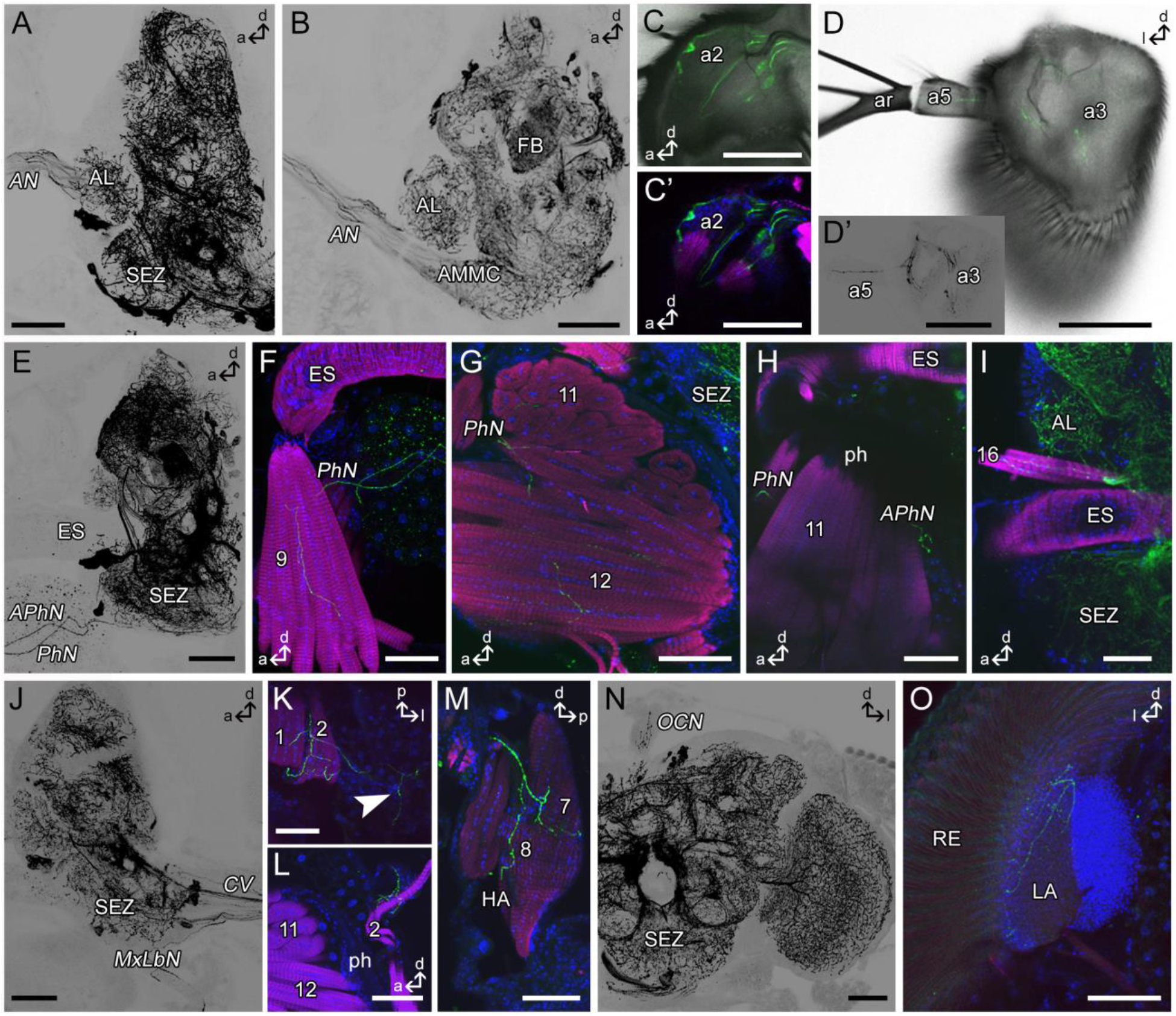
Tdc2-Gal4-positive arborizations in the fly’s head. *Projections of sagittal (A-C, E-J,M) or frontal (D,K,O) or horizontal (L) optical sections visualizing the arborization pattern of Tdc2-Gal4-positive neurons (Tdc2N; black or green) in the head. A-D: Tdc2Ns run through the antennal nerve (AN) and project in antennal segments a2, a3 and a5. Mechanosensory neurons of the Johnston’s organ are visible (C). E-H: Efferent Tdc2Ns of the pharyngeal (PhN) and accessory pharyngeal nerve (APhN). F-G: Cells of the PhN innervate muscles 9, 11 and 12. H: Bouton-like structures of APhN neurons beside the pharynx (ph). I: Innervation of muscle 16. J-M: Tdc2Ns of the maxillary-labial nerve (MxLbN) project along muscle 1 and 2 (K, L) in the haustellum (HA) and seem to innervate muscles 7 and 8 (M). Arborizations from the MxLbN reach the lateral brain (arrowhead in K). N: Tdc2Ns arborize in the ocellar nerve (OCN). O: Ramifications in the lateral lamina (LA) close to the retina (RE). a, anterior; AL, antennal lobe; AMMC, antennal mechanosensory and motor center; ar, arista; d, dorsal; CV, cervical connective; FB, fan-shaped body; l, lateral; p, posterior; SEZ, subesophageal zone. Scale bars = 50 μm.*

The *Tdc2-Gal4*-positive efferent nerves of the subesophageal zone (SEZ) arborize in the rostrum of the proboscis mainly onto muscles (Fig. 2E-M). Cells leaving the brain via the *PhN* innervate muscles 9, 10, 11 and 12 (nomenclature after ^48^; Fig. 2F,G). Cells projecting towards the APhN build bouton like structures beside muscle 11 ventral to the pharynx (ph; Fig. 2H). Cells of the *MxLbN* arborize along muscles 1 and 2 (Fig. 2J-L). It seems as if also muscles 7 and 8 of the haustellum are innervated (Fig. 2M). We only observed this staining in two different specimens. Due to our cutting technique we probably lost these parts of the haustellum frequently. In addition to the innervation of the proboscis muscles we observed arborizations in the ventrolateral head arising from the *MxLbN* (arrowhead Fig. 2K). The ocellar nerve, which connects the ocellar ganglion with the brain, contains fibers arising from the brain (Fig. 2N). The central brain and optic lobes were shown to contain a dense network of OANs/TANs ^37,38^. In addition, we identified arborizations in the distal part of the lamina by Tdc2Ns (Fig. 2O). Muscle 16, which is located dorsal to the esophagus, is innervated via ascending Tdc2Ns from the thorax (Fig. 2I).

### *Tdc2-Gal4*-positive arborizations in the thorax

OANs/TANs form connections between the head and thorax via the *CV* and *NCC* (Fig. 3A). Tdc2Ns running through the *NCC* arborize close to the corpora allata (CA; Fig. 3B) and anterior stomatogastric ganglion, while no staining is visible in the corpora cardiaca (CC; asterisk Fig. 3B). The *CV* connects the brain and VNS and contains many Tdc2Ns (Fig. 3A). All peripheral nerves of the thoracic ganglion seem to contain *Tdc2-Gal4*-positive axons. Most prominent are the paired leg nerves of each thoracic neuromere (*LN1-3* Fig. 3C; ProLN, MesoLN, MetaLN after ^49^), the paired wing *(WN* Fig. 3K; ADMN after ^49^) and posterior dorsal mesothoracic nerve *(PDMN;* Fig. 4A) of the mesothoracic neuromere and paired haltere nerves of the metathoracic neuromere *(HN* Fig. 3N; DMetaN after ^49^). Interestingly, all these nerves, with the exception of *PDMN,* seem to contain efferent Tdc2Ns innervating mainly muscles as well as afferent *Tdc2-Gal4*-positive sensory neurons.

**Fig. 3:**
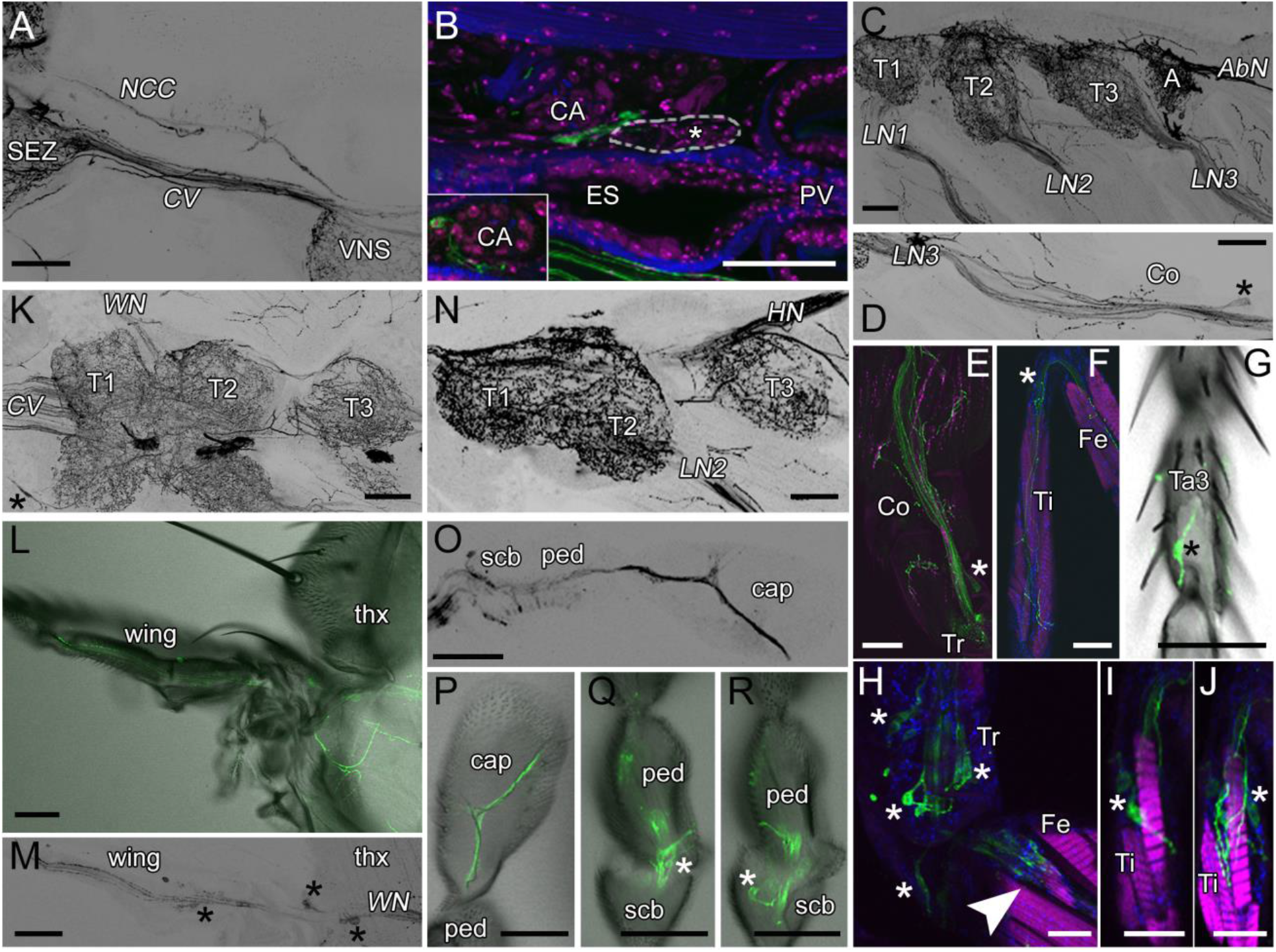
Tdc2-Gal4-positive arborizations in the thorax. *Projections of optical sections visualizing the arborization pattern of Tdc2-Gal4-positive neurons (Tdc2Ns; black or green) in the thorax. A: Tdc2Ns run through the cervical connective (CV) and corpora cardiaca nerve (NCC). B: Tdc2Ns arborize close to the corpora allata (CA) and the anterior stomatogastric ganglion (white-rimmed). C-F: Tdc2Ns project along the legs and innervate leg muscles. G: An afferent sensory neuron in the third segment of the tarsus. H-J: Cell bodies of sensory neurons (asterisks) of the trochanter (Tr; H) and tibia (Ti; I,J). H: Neurons of the chordotonal organ in the femur (Fe; arrowhead). K-M: Tdc2Ns project along the wing nerve (WN). K: Innervation of the thoracic chordotonal organ (asterisk). L,M: Tdc2Ns run along the L1 wing vein. Cell bodies of sensory neurons are visible (asterisks). N-R: Tdc2Ns in the haltere nerve (HN). Tdc2-positive cells project to the distal part of the capitellum (cap). Sensory neurons of the pedicellus (ped) and scabellum (scb) are labeled by Tdc2-Gal4. A-J,N-R: sagittal sections; K: horizontal sections; L,M: frontal sections. A, abdominal segment; Co, coxa; ES, esophagus; Fe, femur; LN, leg nerve; PV, proventriculus; SEZ, subesophageal zone; T1-3, thorax segment1-3; Ta, tarsus; thx, thorax; Ti, tibia; Tr, trochanter. Scale bars: A-G,K-R = 50 μm; H-J = 25 μm.*

**Fig. 4:**
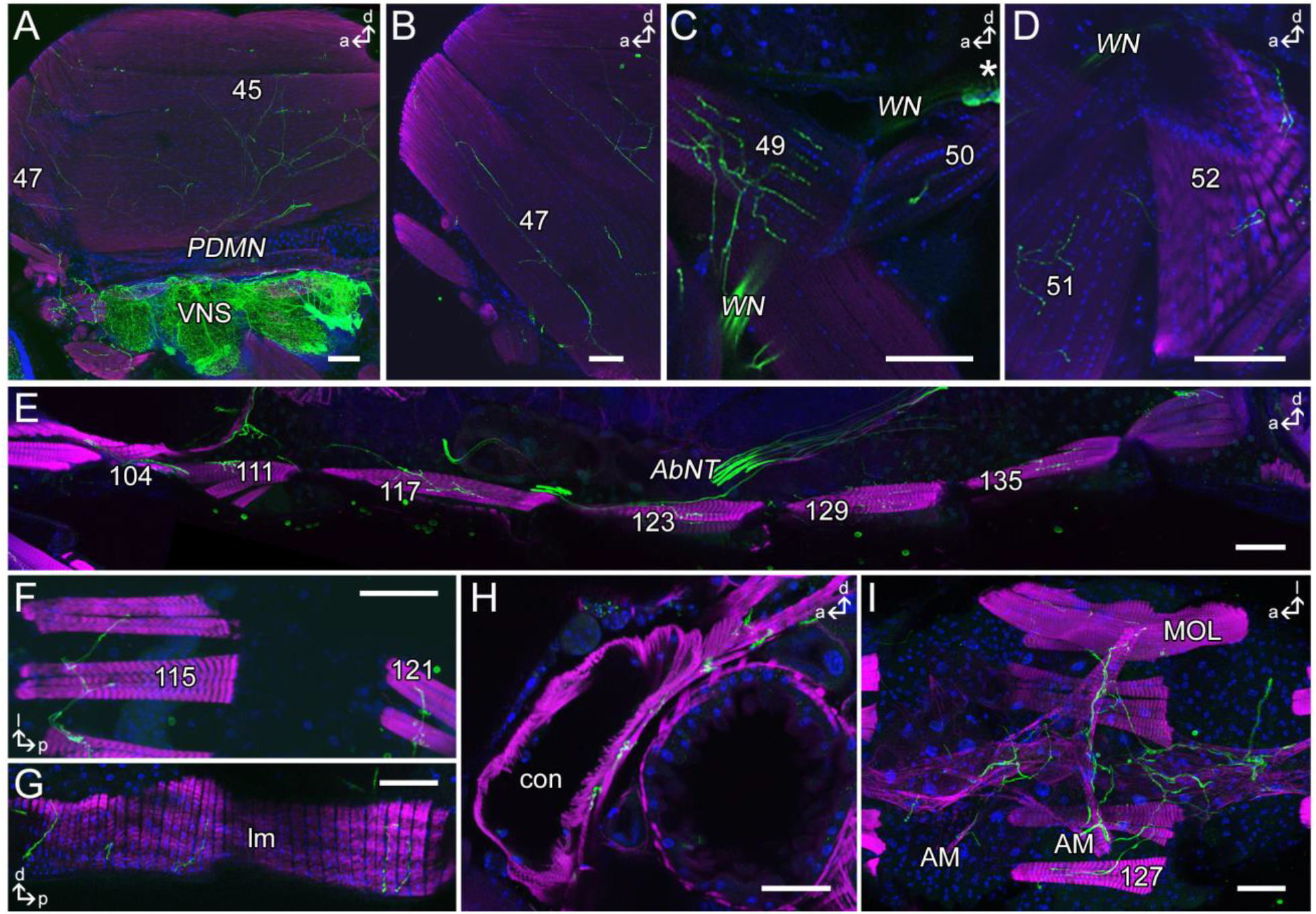
Tdc2-Gal4-positive innervation of skeletal muscles. *Projections of optical sections visualizing the innervation pattern of Tdc2-Gal4-positive neurons (Tdc2Ns; black or green) on skeletal muscles of thorax and abdomen. A-D: Innervation of the indirect flight muscles (A,B) and direct flight muscles (C,D). E-H: Tdc2Ns innervate the ventral (E), dorsal (F) and lateral (G) abdominal body wall muscles. H,I: The longitudinal (H) as well as the alary muscles (I) of the heart are innervated by Tdc2Ns. I: In males arborizations on the muscle of Lawrence (MOL) in segment 5 are visible. A-E,G,H: Sagittal sections; F,I: dorsal sections. a, anterior; AbNT, abdominal nerve trunk; AM, alary muscle; con; conical chamber; d, dorsal; l, lateral; lm, lateral muscles; MOL, muscle of Lawrence; p, posterior; PDMN, posterior dorsal mesothoracic nerve; VNS, ventral nervous system; WN, wing nerve. Scale bars = 50 μm.*

The efferent Tdc2Ns in *LN1-3* arborize on the leg muscles down to the tibia (Fig. 3E,F), while afferent fibers originate from sensory neurons of all leg segments (asterisks Fig. 3D-J), including i.a. mechanosensory neurons of the chordotonal organ of the femur (arrowhead Fig. 3H) and campaniform sensilla of the tarsus (asterisk Fig. 3G). The *WN* contains Tdc2Ns arborizing on indirect and direct flight muscles (Fig. 4B-D) and afferent axons from sensory neurons of the proximal wing (asterisks Fig. 3M, 4C). Moreover, efferent Tdc2Ns running to the PDMN innervate all six longitudinal indirect flight muscles (45a-f; Fig. 4A) and the posterior dorsal-ventral indirect flight muscles (46a-b). Tdc2Ns project along the L1 wing vein (Fig. 3L). Tdc2-positive cells innervating the haltere project to the most distal tip of the capitellum (cap; Fig. 3O,P). Additionally, *Tdc2-Gal4* includes sensory neurons of campaniform sensilla of the pedicellus (ped; Fig. 3Q,R) and scabellum (scb; Fig. 3R). Additionally, it seems that *Tdc2-Gal4* labels sensory neurons of the chordotonal organs of the haltere and wing.

### Tdc2-Gal4-positive arborizations in the abdomen

Tdc2Ns innervate all ventral (111,117,123,129,135; Fig. 4A) and dorsal skeletal muscles (109,115,121,127,133; Fig. 4B) of abdominal segments 2-6 as well as lateral muscles (Fig. 4C). Additionally, *Tdc2-Gal4*-positive ramifications on the male specific “muscle of Lawrence” in segment 5 are visible (Fig. 4E). Beside the body wall muscles, the ventral longitudinal and the alary muscles of the heart are innervated (Fig. 4D,E). Tdc2Ns running along the abdominal nerve trunk (AbNT) innervate the female and male reproductive organs, respectively (Fig. 5). In males, as described before ^41^ the anterior ejaculatory duct, the vas deferens and seminal vesicle are innervated, while the ejaculatory bulb itself is not innervated but its muscles (Fig. 5C-F). The innervations of the female oviducts, uterus and spermathecal duct have been described in previous publications ^36,40–44^.

**Fig. 5:**
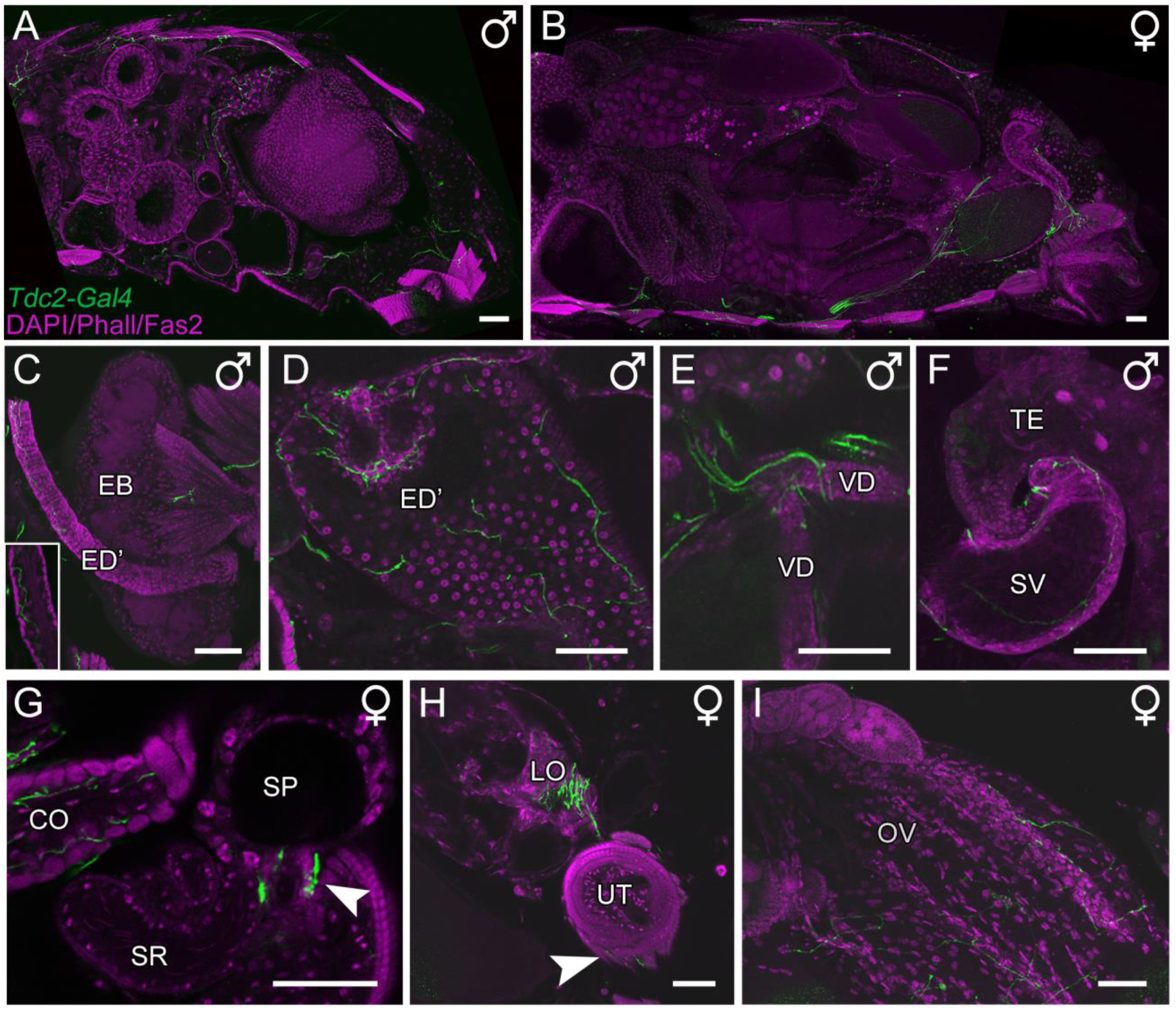
Tdc2-Gal4-positive innervation of the reproductive organs. *Projections of optical sections visualizing the innervation pattern of Tdc2-Gal4-positive neurons (Tdc2Ns; green) of reproductive organs visualized by DAPI, Phalloidin and Fasciclin2 staining (magenta). A-B: Tdc2N arborization pattern in a sagittal section of a male (A) and female (B) abdomen, respectively. C-F: The anterior ejaculatory duct (ED’), muscles of the ejaculatory bulb (EB), the vas deferens (VD) and seminal vesicle (SV) are innervated by Tdc2Ns. G-I: Tdc2Ns arborize onto the common and lateral oviduct (CO, LO), spermathecal duct (arrowhead in G), uterus (UT) muscles (arrowhead in H) and the ovaries (OV). SP, spermatheca; SR, seminal receptacle; TE, testes. Scale bars = 50 μm.*

## Discussion

Here we show a comprehensive description of the innervation of OANs/TANs in the periphery of *Drosophila.* For this, we used the *Tdc2-Gal4* line, allowing Gal4 expression under the control of a regulatory sequence of the tyrosine decarboxylase enzyme ^40^. As this enzyme is essential for the synthesis of TA from tyrosine, the *Tdc2-Gal4*-line labels both TANs and OANs. Within the *Drosophila* brain, *Tdc2-Gal4* labels in total about 137 cells, while additional 39 cells are located in the VNS ^38,39^. The small number of Tdc2Ns lead to arborizations in large parts of the central brain, optic lobes and the thoracic and abdominal ganglion ^37–40^. Based on the profound innervation of Tdc2Ns in the brain and VNS, the variety of behaviors modulated by the OA/TA system including learning and memory, feeding, vision, and sleep, are not surprising. Beyond the brain and VNS, OANs and TANs massively innervate regions within the periphery of the fly. Here, we described arborizations on most skeletal muscles, the antenna, wings, halteres and reproductive system and parts of the circulatory system and stomodaeal ganglion (Fig. 6; Table 1).

**Table 1:** Organs, tissues, visceral and skeletal muscles innervated by *Tdc2-Gal4*-positive neurons (Tdc2Ns).

| Organs, tissues,<br>visceral muscles | Tdc2N<br>staining | Skeletal muscles<br>(after <sup>48</sup>) | Tdc2N<br>staining |
| --- | --- | --- | --- |
| **Digestive system** |  | **Head** |  |
| Esophagus (ES) | o | 1 | x |
| Crop | ? | 2 | x |
| Hindgut | ? | 7 | (x) |
| **Circulatory system** |  | 8 | (x) |
| Corpora allata (CA) | o | 10 | x |
| Ventral longitudinal<br>muscles of heart | x | 11 | x |
| Alary muscles of heart | x | 12 | x |
| **Nervous system** |  | 16 | x |
| Brain | x | **Thorax** |  |
| Ventral nervous system<br>(VNS) | x | **Leg muscles**<br>(after <sup>71</sup>) |  |
| Stomodaeal ganglion | x | Coxa<br>trdm, trlm, trrm | x,x,x |
| Corpora cardiaca (CC) |  | Trochanter<br>fedm, ferm | x,x |
| Ocellar nerve (OCN) | x | Femur<br>ltm2, tilm, tidm, tirm | ?,x,x,x |
| **Sensory organs** |  | Tibia<br>ltm1, talm, tadm, tarm | x,x,x,x |
| Antenna |  | **Flight muscles** |  |
| Pedicle (a2) | x | indirect |  |
| Funiculus (a3) | x | 45,46,47,48 | x,x,x,x |
| Arista (a4,a5) | x | direct |  |
| Wings | x | 49,50,51,52,54,<br>55,56,57,58 | x,x,x,x,x,<br>(x),?,x,? |
| Halteres | x | **Cervical muscles** |  |
| **Male reproductive**<br>**system** |  | D 20,21,22,23,<br>L 24; V 25,26,27 | x,?,x,x,<br>x; x,x,x |
| Muscles of Ejaculatory<br>bulb (EB) | x | **Mesothorax muscles** |  |
| Anterior ejaculatory duct<br>(ED') | x | 59,60,61,62 | x,?,?,x |
| Accessory gland (AG) | ? | **Metathorax muscles** |  |
| Vas deferens (VD) | x | 77,78,79 | ? |
| Seminal vesicle (SV) | x | **Abdomen** |  |
| **Female reproductive**<br>**system** |  | **Segment 1** |  |
| Ovary (OV) | x |  |  |
| Lateral oviduct (LO) | x |  |  |
| Common oviduct (CO) | x |  |  |
| Spermathecal duct (SPd) | x |  |  |
| Spermatheca (SP) |  |  |  |
| Uterus (UT) | x | D 98,99,100,101,102 | x,x,?,x,x |
| Uterus muscles | x | L 103; V 80,81,104 | x; ?,x,x |
| **Sensory organ (SO)** | **Tdc2N cell body** | **Segment 2-6** |  |
| Chordotonal organ (CO) antenna | x | D 109,115,121,127,133 | x,x,x,x,x |
| CO thorax | x | L 110,116,122,128,134 | x,x,x,x,x |
| CO leg | x | V 111,117,123,129,135 | x,x,x,x,x |
| SO leg | x |  |  |
| SO wing | x |  |  |
| SO haltere | x |  |  |

x, staining; o, encircled by staining; (x), not enough samples; ?, ambiguous results or not investigated; D, dorsal; L, lateral; V, ventral

**Fig. 6:**
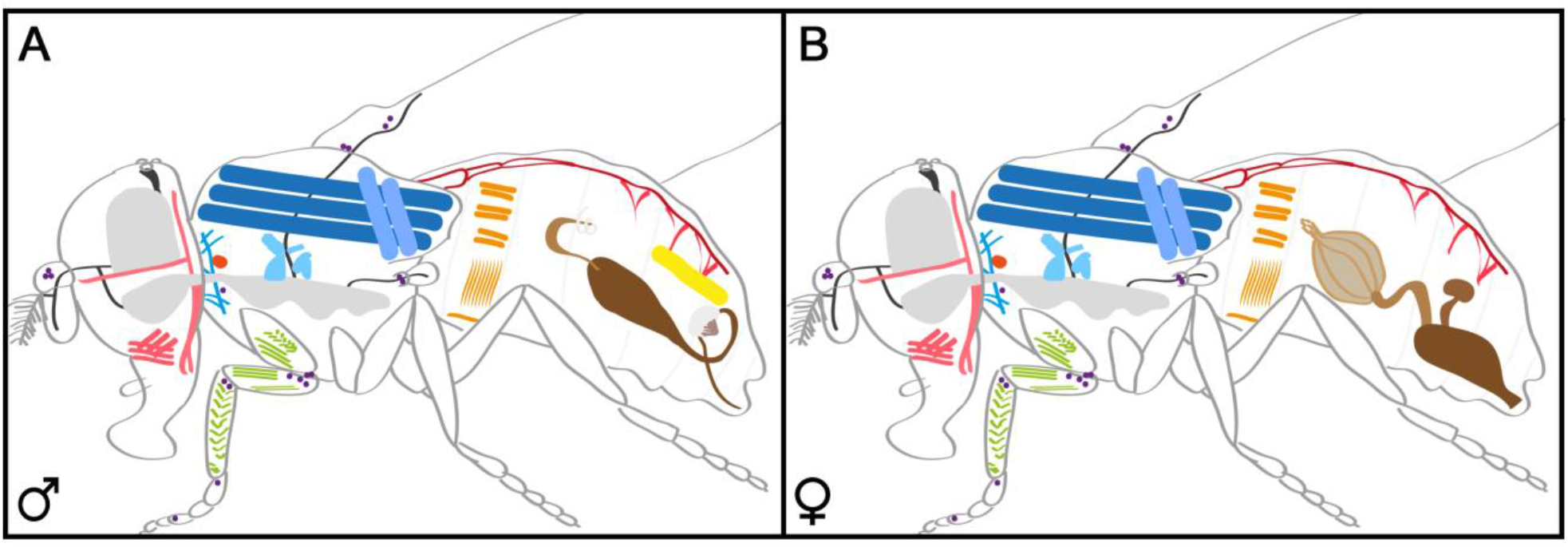
Overview of organs, tissues and skeletal muscles innervated by Tdc2Ns. *Schematic drawing of a male (A) and female (B) fly showing the internal structures innervated by Tdc2Ns: proboscis/head muscles (rose); CNS and VNS (grey); peripheral nerves targeting the antenna, ocelli, wings or halters (dark grey); neck, direct and indirect flight muscles (blue); corpora allata (orange); leg muscles (green); heart and alary muscles (red); abdominal skeletal muscles (ocher); muscle of Lawrence (yellow); reproductive organs (brown). Sensory neurons labeled by the Tdc2-Gal4 line are shown as purple dots.*

Our findings are in line with previous reports focusing on the expression of different OA and TA receptors in the fly ^50,51^. Accordingly, the OA receptor OAMB is expressed in reproductive organs (in both male and female flies) and muscles, which are directly innervated by Tdc2Ns. Additionally, the midgut and trachea contain OA and TA receptors ^50,51^, but do not seem to be innervated by Tdc2Ns, even though axons run in close vicinity to these organs. Likewise, the OA receptor Octβ2R is expressed in the fat body, salivary glands and Malpighian tubules, tissues that seem not to be innervated by Tdc2Ns, while the expression of Octβ1R and Octβ3R is more specific ^50,51^. The three tyramine receptors TyrR, TyrRII and TyrRIII show a broad expression in the periphery. Interestingly, TyrR seems to be the only receptor expressed in the heart, suggesting that only TA modulates heart function ^50^. Contrary, OA has a modulatory effect on the heart of other insect species including honeybees, olive fruit flies and cockroaches ^52^. This is also in line with a previous report providing evidence that OA modulates the heart rate of the *Drosophila* fly and pupa, but not the larva ^53^.

OA-dependent modulation of organs and tissues is mainly elicited through muscle action, especially in terms of its impact on the “fight or flight” response. In line with this, we observed *Tdc2-Gal4*-positive arborizations on nearly all skeletal muscles and many visceral muscles (Table 1). In both *Drosophila* and desert locusts OA and TA is expressed in type II terminals of skeletal muscles ^45^. OA has an excitatory effect on *Drosophila* flight muscles, while TA was shown to inhibit excitatory junction potentials, and thereby reduce muscle contractions and locomotion at least in the larva ^6,8,54,55^. In addition, flies lacking OA show severe deficits in flight initiation and maintenance ^10,45^. Interestingly, in an antagonistic effect to serotonin, OA reduces crop muscle activity presumably via Octβ1R, suggesting that OA has different effects on muscle activity dependent on the type of muscle ^50,56^. However, our data do not provide any evidence of a direct innervation of Tdc2Ns of the crop, even though many fibers run in close vicinity, suggesting that OA might target the crop by volume transmission.

Furthermore, OA modulates ovulation and fertilization in insects ^57–61^. Flies lacking OA display a severe egg-laying phenotype. Remarkably, within the female reproductive organ two different OA receptors, OAMB and Octβ2R, are necessary. Again, OA has a strong impact on muscle activity within the reproductive system. Octβ2R is expressed in the visceral oviduct muscle and elicits muscle relaxation through an increase of intracellular cAMP levels ^57^. Such an OA-dependent modulation appears to be conserved as OA is found in dorsal unpaired median neurons of locusts innervating oviduct muscles through the oviducal nerve ^62^. However, our data suggest that OA-positive fibers not only innervate oviduct muscles, but also enter the organs themselves. The OAMB receptor is expressed in ephitelial cells inducing fluid secretion through increasing intracellular Ca^2+^ levels ^57^. Thus, OA affects different processes within the female reproductive organ due to the expression of different receptors and their coupled signaling pathways, which may be a general mechanism of the OA/TA system to fulfill an extensive modulatory function ^63^.

OA does not exclusively modulate muscle activity, but also sensory neurons of external tissues like the antenna, halteres and wings. OA has also been shown to increase the spontaneous activity of olfactory receptor neurons (ORN) ^64,65^. The modulation of ORNs allows OA to modulate the innate response to attractive stimuli like fruit odors or pheromones ^66,67^. Further, this modulation helps nestmate recognition in ants ^68^. In addition to *Tdc2-Gal4*-positive arborizations in the funiculus, we found *Tdc2-Gal4-* positive sensory neurons in the Johnston’s organ, a chordotonal organ sensitive to mechanosensory stimuli and thus important for hearing in insects. In mosquitos, OA modulates auditory frequency tuning and thereby affects mating behavior ^69^. In locusts, OA similarly modulates the response of chordotonal neurons in the legs to encode proprioceptive information ^70^. Our data suggest that chordotonal neurons in the leg, wings, halteres and thorax are included in the *Tdc2-Gal4* line suggesting a conserved modulatory role of OA/TA for insect proprioception.

Taken together, our study suggest that the OA/TA system massively modulates various organs and tissues in the periphery of *Drosophila.* Through distinct receptors and coupled signaling pathways OANs/TANs mainly induce “fight or flight” responses by modulating muscle activity, proprioception, and heart rate. As a result, the innervation pattern in the periphery supports the idea that the OA/TA system is crucial for insects to switch from a dormant to an excited state, by a positive modulation of muscle activity, heart rate and energy supply, and a simultaneous negative modulation of physiological processes like e.g. sleep.

## Methods

### Fly strains and fly rearing

All flies were cultured according to standard methods. In short, vials were kept under constant conditions with 25°C and 60% humidity in a 12:12 light:dark cycle. Flies carrying the *Tdc2-Gal4* (^40^, Bloomington Stock Center) and *10xUAS-IVS-myrGFP* (^72^, Bloomington Stock Center) constructs were used for immunohistochemistry. To control for an unspecific expression of the UAS construct, we stained *10xUAS-IVS-myrGFP* alone. No GFP staining was detected.

### Immunocytochemistry

To visualize the arborisations of *Tdc2-Gal4-positive* neurons in the periphery whole body sections, as well as sections of the head, thorax and abdomen, were performed, respectively (see ^73^). In short, the cuticle of 4 to 7 days old flies was opened in phosphate buffered saline (PBS, 0.1M) to ensure that the fixative is able to penetrate into the tissue. Whole flies were fixated with 4% paraformaldehyde in PBS for two hours at room temperature and afterwards washed three times with PBS. Subsequently flies were embedded in hot 7% Agarose low EEO (A2114; AppliChem). After hardening, the flies were cut with a vibratome (Leica VT1000S) into 80-100 μm sections. Staining of the sections was continued after washing in PBS containing 0.3% Triton-X100 (PBT) and blocking in 5% normal goat serum in PBT. Rabbit anti-GFP (A6455, Molecular Probes) in combination with mouse anti-Synapsin (3C11; ^74^; 1:50) or mouse anti-Bruchpilot (nc82; ^75^) and mouse anti-Fasciclin 2 (1D4; DSHB; 1:100) were used as primary antibodies. After one night at 4°C the specimens were washed six times in PBT and incubated in secondary antibody solution for a subsequent night at 4°C. As secondary antibodies goat anti-rabbit Alexa488 (Molecular Probes; 1:200) and goat anti-mouse DyLight649 (Jackson ImmunoResearch; 1:200) were used. 4’,6-Diamidino-2-phenylindol Dihydrochlorid (DAPI; Sigma-Aldrich; 1:1000) and Alexa Fluor 633 Phalloidin (Molecular Probes; 1:400) were used to visualize DNA and actin, respectively.

### Confocal microscopy and data processing

Confocal images were taken with a Leica TCS SP8 microscope (Leica Microsystems, Germany) with a 20x high aperture objective. Labelled specimens were scanned with a step size of 1.0 μm to 1.5 μm. Image processing and alignment was performed using Fiji (^76^), Amira 5.3 (Visage Imaging, Berlin, Germany and Adobe Photoshop CS6 (Adobe Systems, USA).

## Acknowledgements

We thank the Bloomington stock center for flies, Claudia Groh and the Development Studies Hybridoma Bank for antibodies. We thank Christian Wegener and Charlotte Förster for their support and fruitful discussions, and Christian Wegener for comments on the manuscript.

This research was supported by a grant from the German Excellence Initiative to the Graduate School of Life Sciences, University of Würzburg (to M.S.); SCIENTIA fellowship “Bayerische Gleichstellungsförderung: Programm zur Realisierung der Chancengleichheit für Frauen in Forschung und Lehre” (to M.S.), and by the Deutsche Forschungsgemeinschaft (PA1979/2-1 to D.P., EL784/1-1 to B.eJ.).

D.P. and M.S. conceived and designed the experiments. D.P., C.B., F.F., B.eJ. and M.S. performed the experiments. M.S. analyzed the data. D.P. and M.S. wrote the manuscript and M.S. prepared the figures. All authors reviewed the manuscript.

